# Decoy and antisense RNA oligonucleotides targeting the EGFR exon 3 splice site suppress proliferation of A172 glioblastoma cells

**DOI:** 10.64898/2026.10.25.761620

**Authors:** Yumika Nishi, Yuta Horita, Hiroyoshi Harada, Yusuke Murata, Masayoshi Mori, Kenji Ohe

**Author notes:** Corresponding author: Kenji Ohe.

## Abstract

Short RNA oligonucleotides directed at the 3’ splice site region of *EGFR* exon 3 reduced the proliferation of A172 glioblastoma cells, while the same oligonucleotides had no detectable effect on A549 lung adenocarcinoma cells.

Glioblastoma carries *EGFR* amplification in roughly a third of IDH-wildtype cases, and the A172 line harbours a tandem duplication in the *EGFR* tyrosine kinase domain. Because exon definition in human genes depends on RNA-binding proteins that engage exonic sequence near splice sites, we used the SpliceAid database to identify candidate RNA-binding proteins at *EGFR* exons 2 and 3. Y-box binding protein 1 (YB-1, encoded by *YBX1*), reported to drive glioblastoma growth through its RNA-binding function, scored highly immediately adjacent to the exon 3 3’ splice site.

We designed four 2’-modified RNA oligonucleotides carrying a nuclear localisation signal: two antisense oligonucleotides against the exon 3 region (asRNA1, spanning the splice site; asRNA2, downstream of it), a decoy complementary to the asRNA2 target (dRNA2), and a decoy carrying the YB-1 consensus binding sequence (dYB-1). In A172 cells, all four reduced EGFR protein in a single Western blot experiment, and asRNA1, asRNA2 and dRNA2 reduced proliferation relative to scrambled RNA at days 1–3; dYB-1 produced a comparable reduction at day 3. None of these effects was seen in A549 cells. YB-1 protein level and subcellular distribution were unchanged by the oligonucleotides, arguing against an indirect route through altered YB-1 abundance.

These observations are preliminary. Every experiment reported here was performed once: the proliferation data come from single plates with three wells per group, analysed by Dunnett’s test against the scrambled-RNA control, and the Western blot, RT-PCR and immunocytochemistry panels are single runs presented as representative observations. Direct binding of YB-1 to the targeted *EGFR* sequence has not been demonstrated. We post this work as a preprint so that the design strategy and the initial observations are available while these questions remain open.

## Introduction

Glioblastoma remains close to incurable. It accounts for about 11% of primary brain tumours, peaks in the fifth and sixth decades, and is classified as WHO grade 4 [1]. Even with maximal surgery, radiotherapy and temozolomide, median survival is 14.6 months and five-year survival is 10.1% [2]. Chemotherapy was long considered ineffective [3] until temozolomide, which crosses the blood–brain barrier, entered the standard regimen [4]; recurrence nonetheless occurs within months to a year of initial treatment in most patients [5]. Photodynamic therapy and tumour-treating fields have since been approved in Japan [6], but age, preoperative performance status, extent of resection and postoperative performance status still dominate prognosis.

The 2021 WHO classification of central nervous system tumours moved diagnosis onto molecular ground, with *IDH, TP53, TERT, EGFR, CDKN2A/B, H3 K27M* and *BRAF* status and 1p/19q codeletion defining entities [7]. Among IDH-wildtype glioblastoma, which is about 90% of cases, *EGFR* amplification is present in roughly 35% [8]. The A172 line used here carries a tandem duplication of the EGFR tyrosine kinase and calcium internalisation domains, conferring ligand-independent signalling [9]. EGFR is therefore a rational target in this background.

Our approach was to attack *EGFR* expression at the level of RNA processing rather than at the protein. Human introns are long relative to exons, so splice sites are recognised across the exon — exon definition — and RNA-binding proteins engaging exonic sequence near the intron boundary are central to that recognition [10]. Targeting a downstream exon risks leaving upstream translation intact and producing a truncated receptor, so we focused on exons 2 and 3, close to the 5’ end of the 26-exon gene.

Searching the SpliceAid database for RNA-binding proteins at these boundaries returned YB-1 (gene symbol *YBX1*) with a high score immediately adjacent to the exon 3 3’ splice site. YB-1 is a multifunctional DNA- and RNA-binding protein acting in transcription, splicing, mRNA stability and translational control, and it has been reported to promote glioblastoma growth through a YB-1/CCT4/mLST8/mTOR axis, with a YB-1 decoy RNA extending survival in a mouse model [11]. YB-1 has also been reported to enhance EGFR signalling through translational mechanisms [14]. This convergence — a predicted YB-1 site at an exon we wished to disrupt, in a protein already implicated in glioblastoma — motivated the oligonucleotide set described below.

## Materials and Methods

### Cell culture

A172 (glioblastoma) and A549 (lung adenocarcinoma, EGFR wild type) cells were maintained in DMEM (Nacalai Tesque) with 10% FBS (Japan Bio Serum) at 37 °C under 5% CO_2_.

### RNA oligonucleotides

All oligonucleotides carried a 3’ nuclear localisation signal (underlined), as described for microRNAs [10].

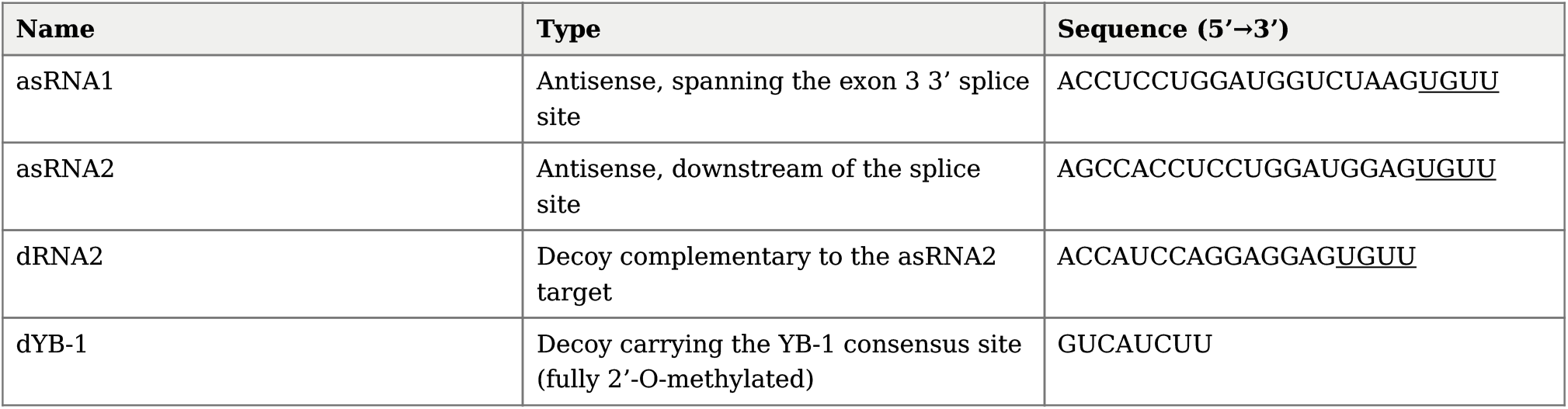

asRNA1 overlaps the splice site and was used as a positive control for disrupted exon definition; asRNA2 tests the contribution of the downstream sequence alone; dRNA2 and dYB-1 act as decoys for the RNA-binding proteins engaging those sequences.

### Transfection

Cells were seeded in six-well plates and transfected at approximately 80% confluence with 3 µg RNA oligonucleotide and 3 µL Lipofectamine 3000, complexed for 20 min at room temperature before addition.

### Proliferation assay

Cells were seeded at 1,000 per well in 96-well plates, transfected, and cultured for 24, 48 and 72 h. Viable cells were stained with a Crystal Violet Assay Kit (Abcam) and absorbance read at 595 nm (Tecan); values are expressed relative to the 0 h reading. Each assay was performed once, with three wells per treatment group on a single plate; the three values per group therefore represent wells within one experiment rather than independent experiments. Data are shown as mean ± SD of those three wells. Because each treatment group is compared against the same scrambled-RNA control, differences were assessed by Dunnett’s test rather than by repeated two-sample t-tests. The A172 and A549 panels of Fig. 6 derive from separate experiments performed on different days.

### RT-PCR

Total RNA was extracted with Trizol (Invitrogen) and chloroform, reverse transcribed from 1 µg with ReverTra Ace (Toyobo), and amplified with Ex Taq (Takara). Primers: EGFR forward TCAGCCTCCAGAGGATGTTC, reverse GTGTTGAGGGCAATGAGGAC; ACTB forward TCACCCACACTGTGCCCATCTACGA, reverse CAGCGGAACCGCTCATTGCCAATGG.

### Western blotting

Protein was recovered by acetone precipitation from the interphase of the Trizol extraction, dissolved in urea sample buffer (10 mM Tris pH 7.5, 25 mM KCl, 5 mM MgCl_2_, 6 M urea) and sonicated. Approximately 20 µg per lane was separated by SDS-PAGE, transferred to PVDF, blocked with Blocking One (Nacalai Tesque) for 10 min at 4 °C, incubated with primary antibody (1:1,000) for 16 h at 4 °C and secondary antibody (1:5,000) for 1 h at room temperature, and detected with Chemi-Lumi One L (Nacalai Tesque). Antibodies: anti-EGFR (4267T, Cell Signaling), anti-YBX1 (PAC609Hu01, Cloud-Clone), anti-ACTB (A00702, CGCS), anti-mouse IgG HRP (7076S, Cell Signaling).

### Immunocytochemistry

A172 cells on chamber slides (Iwaki 5732-008) were transfected, fixed in 4% paraformaldehyde for 15 min, permeabilised with Triton X-100 for 15 min, blocked in 5% BSA / 0.1% Tween-20 for 60 min, incubated with anti-YB-1 overnight at 4 °C, then with Alexa Fluor 568 goat anti-rabbit IgG (1:500, Thermo Fisher A11011) for 1 h protected from light, and counterstained with Hoechst 33258 (1:1,000, 10 min). Imaging used a BZ-X710 microscope (Keyence). The NMD inhibitor was HY-111374 (MedChemExpress).

### In silico analysis

RNA-binding protein sites flanking *EGFR* exon–intron boundaries were predicted with SpliceAid (http://www.introni.it/splicing.html).

### Replication

Every experiment reported here was performed once. For the proliferation assay this means one plate per experiment with three wells per group; for the Western blot, RT-PCR and immunocytochemistry it means a single run, presented as a representative observation without densitometric quantification or statistical testing. Independent replication has not been established for any of the results in this report.

## Results

### Prediction of RNA-binding protein sites near EGFR exons 2 and 3

The *EGFR* gene comprises 26 exons, with a long intron between exons 1 and 2 (Fig. 1). SpliceAid analysis of the sequences flanking the exon 2 and exon 3 splice sites returned several RNA-binding proteins (Fig. 2). YB-1 scored highly at the 5’ splice site region of exon 2 and, more strikingly, immediately adjacent to the 3’ splice site of exon 3 (Fig. 3).

**Fig. 1.**
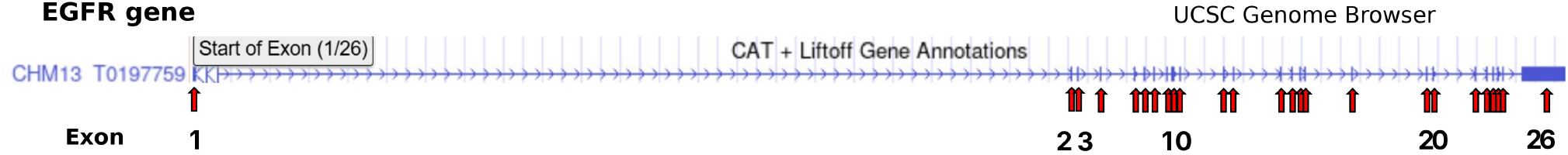
Exon–intron structure of the *EGFR* gene. The gene comprises 26 exons, with a long intron between exons 1 and 2. Image generated with the UCSC Genome Browser (https://genome.ucsc.edu).

**Fig. 2.**
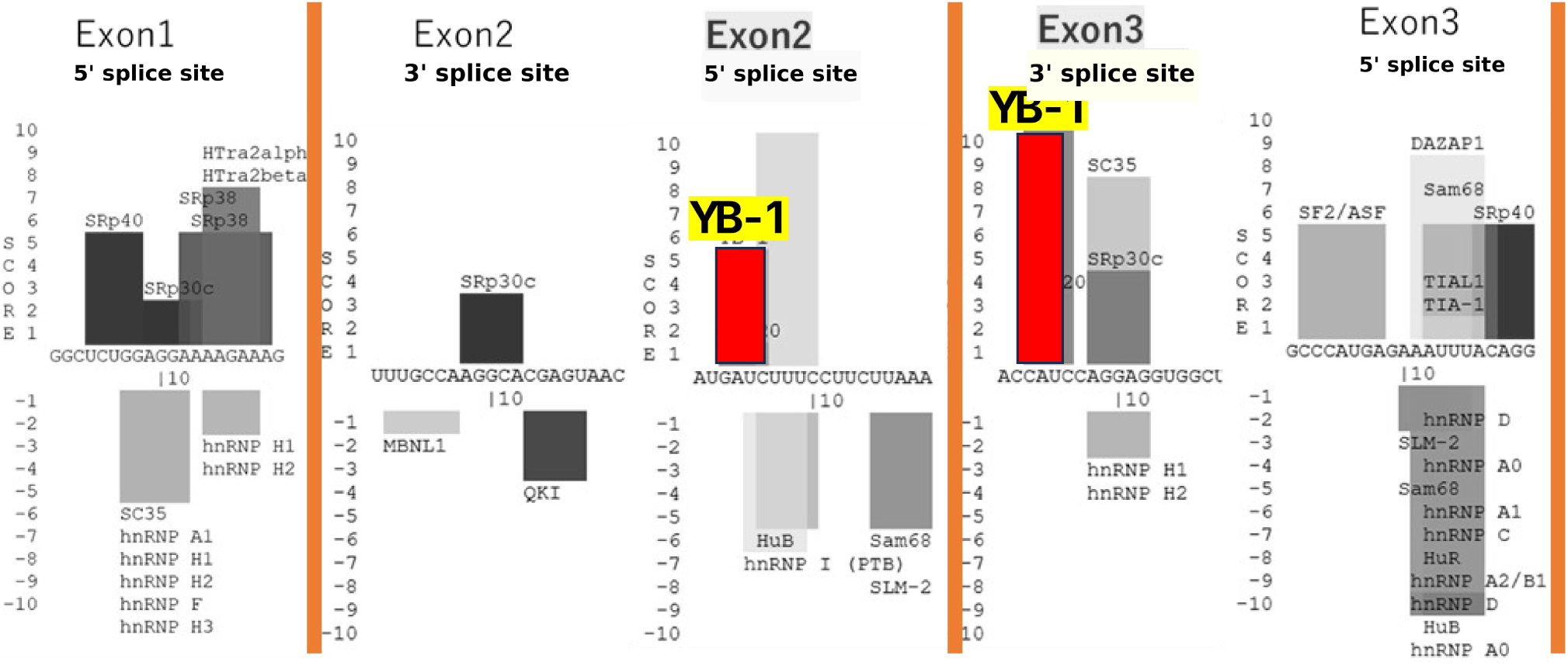
RNA-binding proteins predicted at the *EGFR* exon 2 and exon 3 splice sites. Output of the SpliceAid database (http://www.introni.it/splicing.html) for the sequences flanking the 5’ and 3’ splice sites of exons 1–3. Bars above the sequence indicate proteins predicted to bind the sense strand; bars below, the antisense strand. YB-1 (highlighted) scores highly at the exon 2 5’ splice site region and immediately adjacent to the exon 3 3’ splice site.

**Fig. 3.**
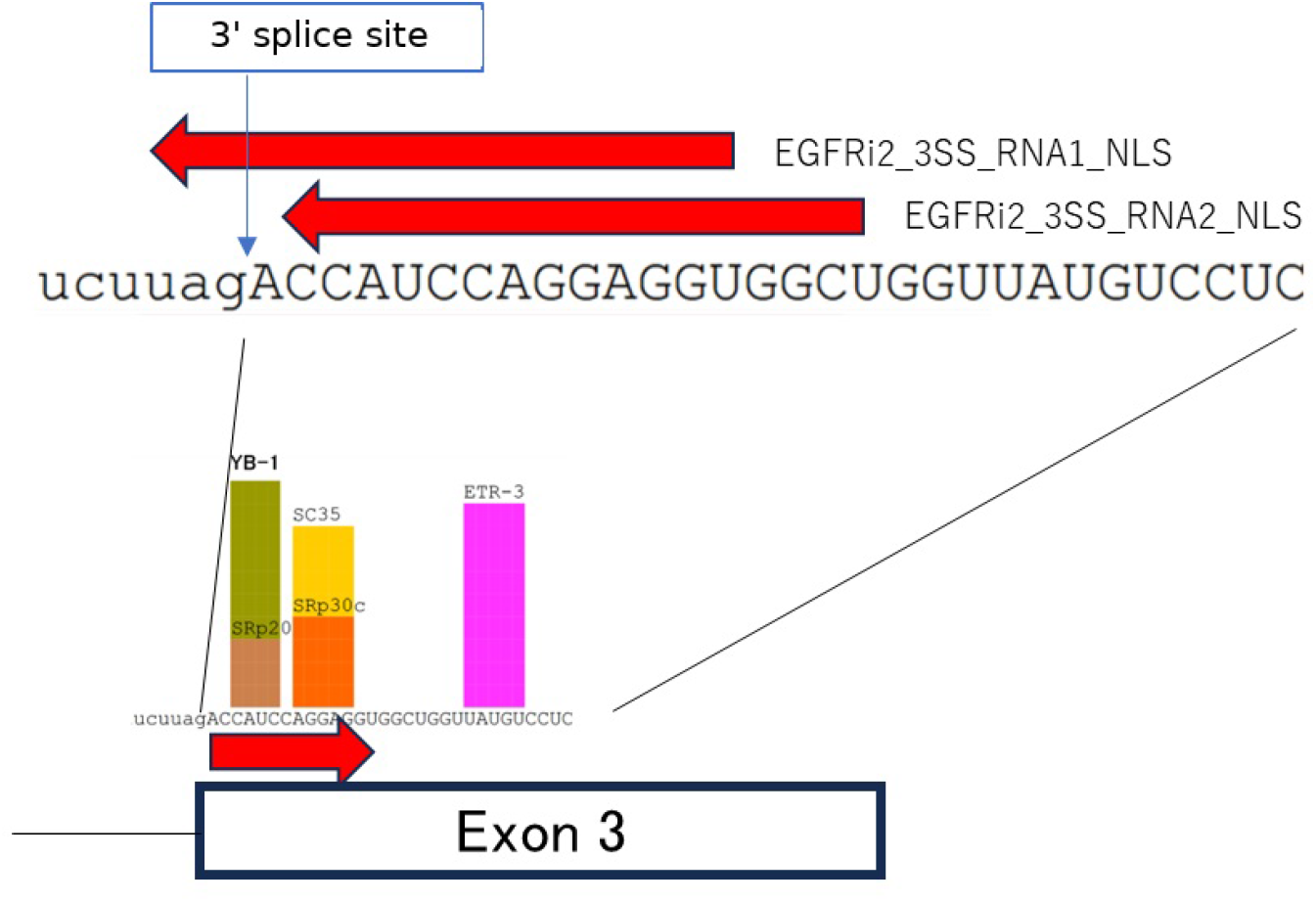
Design of the RNA oligonucleotides against the exon 3 3’ splice site region. The exon 3 3’ splice site and the adjacent exonic sequence are shown with the predicted binding proteins (SpliceAid, http://www.introni.it/splicing.html). Arrows mark the regions targeted by asRNA1 (EGFRi2_3SS_RNA1_NLS), which spans the splice site, and asRNA2 (EGFRi2_3SS_RNA2_NLS), which lies downstream of it.

We designed two antisense oligonucleotides against this region: asRNA1, which overlaps the splice site itself and would be expected to block splicing directly, and asRNA2, which lies downstream of it. Two decoys were also prepared: dRNA2, complementary to the asRNA2 target and therefore expected to sequester the RNA-binding proteins engaging that sequence, and dYB-1, carrying the YB-1 consensus binding sequence.

### Effect on EGFR protein and mRNA

In a single Western blot experiment, all four oligonucleotides lowered the EGFR/ACTB signal ratio in A172 cells relative to scrambled RNA (1.6 for scrambled; 0.8, 0.3, 0.2 and 0.2 for asRNA1, asRNA2, dRNA2 and dYB-1), whereas the corresponding A549 lanes showed no consistent reduction (Fig. 4). The reduction in A172 cells was larger for asRNA2, dRNA2 and dYB-1 than for asRNA1, the splice-site-overlapping positive control. These are single-experiment values and are reported without statistical testing.

**Fig. 4.**
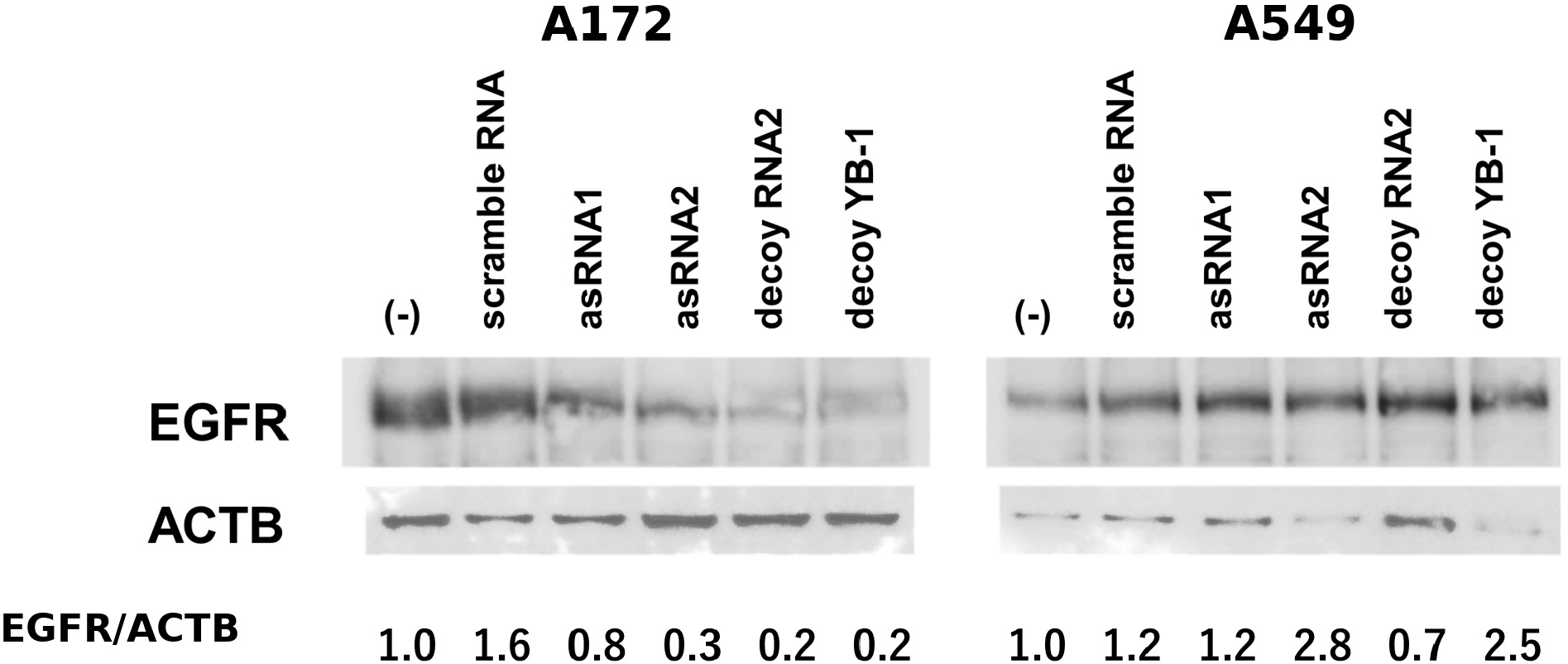
Effect of the RNA oligonucleotides on EGFR protein. Western blot of A172 and A549 cells 48 h after transfection. EGFR/ACTB signal ratios are given below each lane, normalised to the untransfected control. Single experiment; the ratios are single measurements and were not tested statistically.

Semi-quantitative RT-PCR at 48 h showed reduced EGFR signal in A172 cells transfected with asRNA1 or decoy RNA1, but the ACTB control signal was reduced in the same lanes (Fig. 5), so these panels do not establish a specific change in EGFR mRNA. Cell morphology appeared unaffected by light microscopy. ACTB transcript levels can vary with differentiation and metabolic state, and normalisation against ribosomal protein genes such as *RPL19* and *RPL29* has been recommended in such settings [12]. The corresponding A549 lanes showed neither change.

**Fig. 5.**
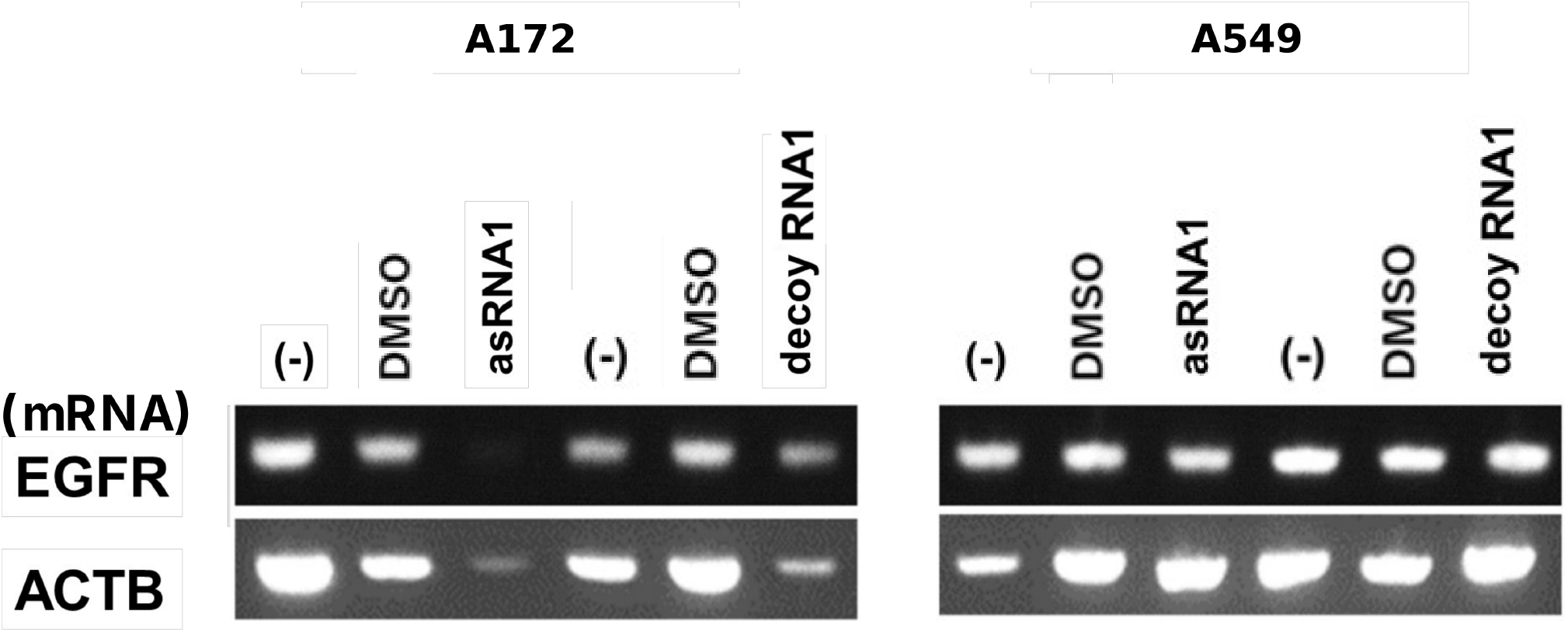
Effect of the RNA oligonucleotides on EGFR mRNA. Semi-quantitative RT-PCR of A172 and A549 cells 48 h after transfection. The ACTB signal falls together with the EGFR signal in the A172 lanes, so this panel does not establish a specific change in EGFR transcript. Single experiment.

### Effect on proliferation

In A172 cells, asRNA2 and dRNA2 lowered proliferation relative to scrambled RNA at every time point, and asRNA1 did so at day 2 (Fig. 6A; Dunnett’s test against scrambled RNA, three wells per group). At day 1 the values relative to the day 0 reading were 2.64 for scrambled RNA, 2.21 for asRNA1 (p = 0.093), 1.52 for asRNA2 (p = 0.0004) and 1.29 for dRNA2 (p = 0.0001). At day 2 they were 2.86, 2.10 (p = 0.012), 1.29 (p = 0.0001) and 1.19 (p = 0.0001), and at day 3 5.70, 3.21 (p = 0.062), 2.60 (p = 0.023) and 1.79 (p = 0.0067).

**Fig. 6.**
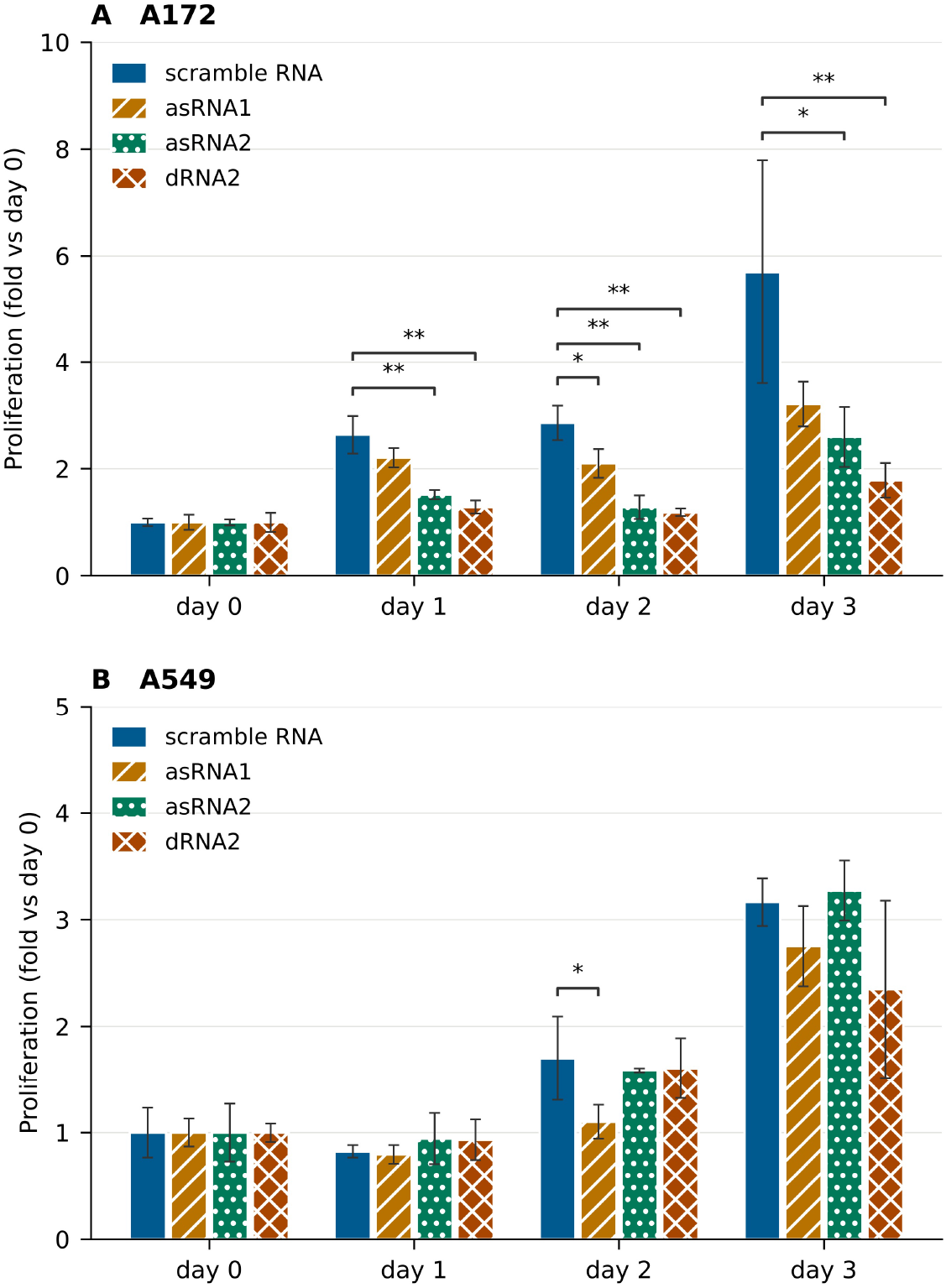
Effect of the RNA oligonucleotides on proliferation. (A) A172 and (B) A549 cells, stained with crystal violet and read at 595 nm, expressed relative to the day 0 reading. Bars are mean ± SD of three wells on a single plate; panels A and B are separate experiments performed on different days. *p < 0.05, **p < 0.01 versus scrambled RNA (Dunnett’s test).

In A549 cells the same oligonucleotides produced no comparable reduction (Fig. 6B). At day 3 the values were 3.17 for scrambled RNA, 2.75 for asRNA1 (p = 0.62), 3.27 for asRNA2 (p = 0.99) and 2.35 for dRNA2 (p = 0.17); the only nominally significant difference across the time course was asRNA1 at day 2 (1.10 versus 1.70, p = 0.050).

In a separate experiment including the YB-1 decoy, dYB-1 lowered A172 proliferation at day 3 (2.06 versus 5.51 for scrambled RNA, p = 0.016; Fig. 7B), with no difference at day 1 (p = 0.16) or day 2 (p = 0.39). In that experiment asRNA2 and dRNA2 did not differ from scrambled RNA at day 3 (p = 0.64 and p = 0.999).

**Fig. 7.**
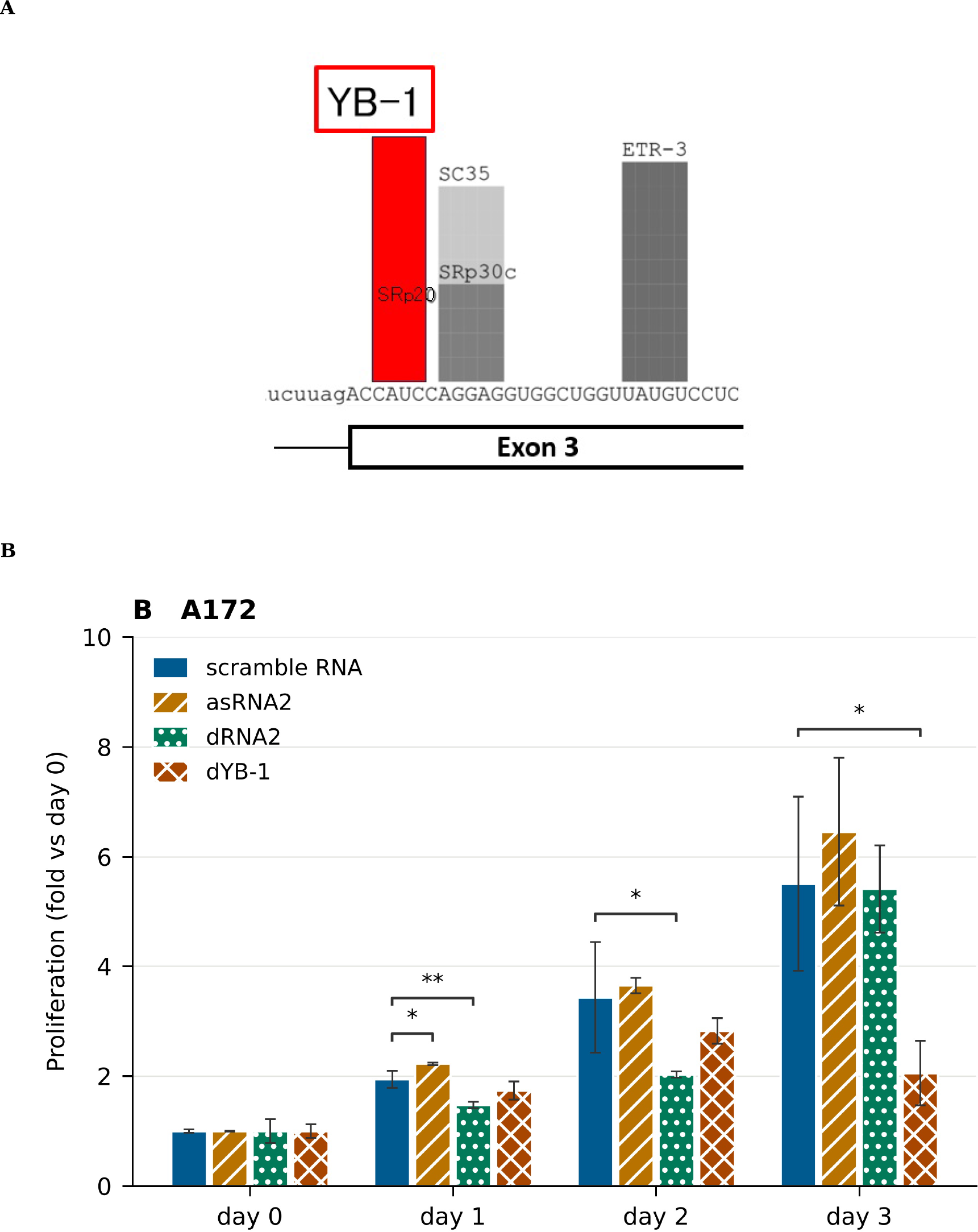
YB-1 at the exon 3 3’ splice site and the effect of the YB-1 decoy. (A) SpliceAid prediction for the exon 3 3’ splice site region (http://www.introni.it/splicing.html); YB-1 is highlighted. (B) Proliferation of A172 cells transfected with scrambled RNA, asRNA2, dRNA2 or dYB-1, from an experiment separate from that in Fig. 6. Bars are mean ± SD of three wells on a single plate. *p < 0.05, **p < 0.01 versus scrambled RNA (Dunnett’s test).

### YB-1 level and localisation

To ask whether the oligonucleotides act indirectly by changing YB-1 abundance or distribution, we examined YB-1 protein, the effect of an NMD inhibitor, and YB-1 immunostaining (Fig. 8). YB-1 protein levels in A172 and A549 cells were not markedly altered by any of the oligonucleotides (Fig. 8A), and YB-1 immunostaining showed no evident redistribution between nucleus and cytoplasm with or without NMD inhibition (Fig. 8C). An RT-PCR panel spanning exons 2–4 resolved products of 470 bp (exons 2, 3, 4) and 286 bp (exons 2, 4), the latter corresponding to exon 3 skipping (Fig. 8B). These panels are single experiments and were not quantified.

**Fig. 8.**
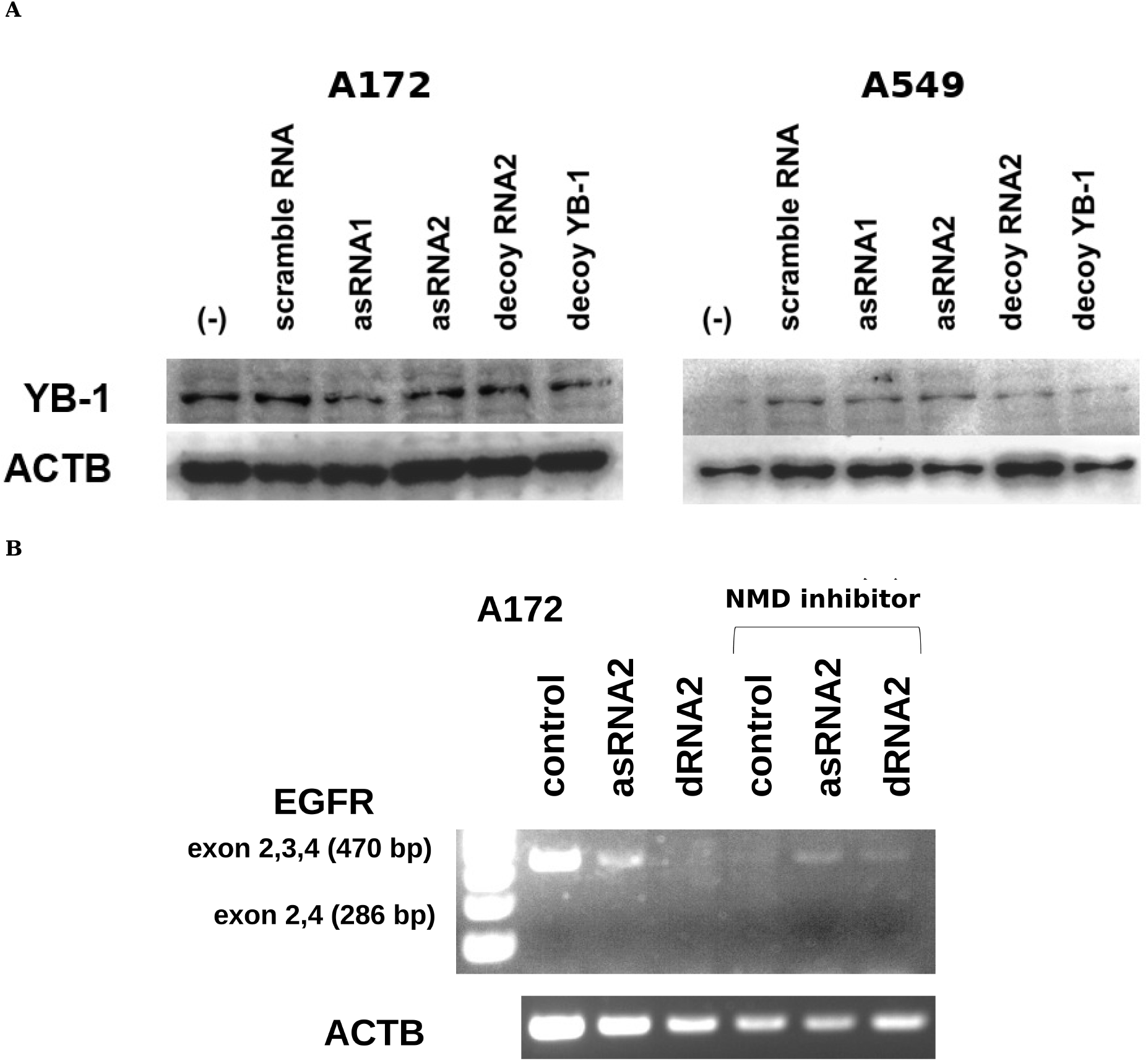
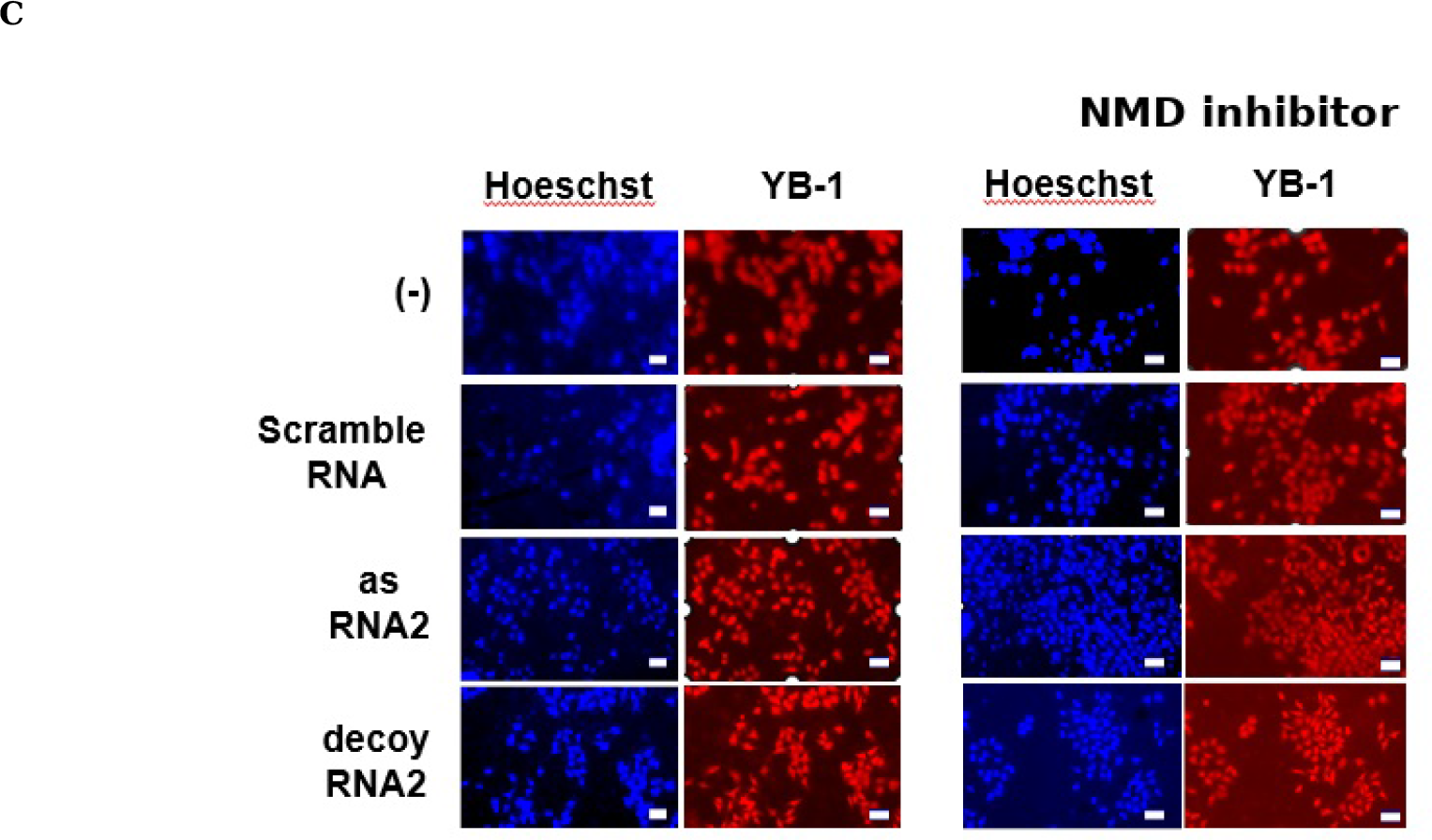
YB-1 protein level, exon 3 skipping and YB-1 localisation. (A) Western blot of YB-1 in A172 and A549 cells 48 h after transfection. (B) RT-PCR across exons 2–4 of *EGFR* in A172 cells with and without the NMD inhibitor HY-111374, resolving products of 470 bp (exons 2, 3, 4) and 286 bp (exons 2, 4; exon 3 skipped). (C) YB-1 immunostaining (red) with Hoechst 33258 counterstain (blue) in A172 cells, with and without NMD inhibition. Scale bars, 50 µm. Each panel is a single experiment and was not quantified.

## Discussion

RNA oligonucleotides directed at the exon 3 3’ splice site region of *EGFR* reduced A172 proliferation in the experiments reported here, and the two decoys — one complementary to the targeted sequence, one carrying the YB-1 consensus site — were as effective as the antisense oligonucleotides. That a decoy for an RNA-binding protein reproduces the effect of an antisense oligonucleotide against the same region is consistent with the sequence acting through protein binding rather than through steric block alone, though we have not demonstrated this directly.

YB-1 is a plausible candidate for that protein. It acts in transcription, splicing, mRNA stability and translational control, and through its DNA-binding function regulates P-glycoprotein (*MDR1*) and thereby drug resistance; its overexpression in glioblastoma is associated with progression and treatment resistance [13]. Wang and colleagues reported that YB-1 binds the 5’UTR of *CCT4* mRNA, promotes its translation, stabilises mLST8 and activates mTORC1 and mTORC2, and that a YB-1 decoy RNA suppressed tumour growth and extended survival in mice [11]. EGFR activation sits upstream of that cascade, and YB-1 has separately been reported to promote translation of developmental transcription factors including Snail1 and to enhance EGFR signalling [14]. Our observations are compatible with a route in which YB-1 contributes to EGFR expression at the level of RNA processing, but the experiments here do not distinguish this from alternatives.

The exon 2–4 RT-PCR panel (Fig. 8B) resolved a product consistent with exon 3 skipping. Quantifying that product under each oligonucleotide would test the exon-definition mechanism directly and is, in our view, the most informative next experiment.

The absence of any effect in A549 cells has at least three explanations we cannot separate: lower transfection efficiency, reduced accessibility of the exon 3 target in that transcript, and genuine dependence of the effect on the *EGFR* tandem duplication present in A172. Measuring transfection efficiency, for instance by electroporation with a labelled oligonucleotide and TaqMan quantification, would distinguish the first from the others.

### Limitations

Every experiment reported here was performed once, and independent replication has not been established. In the proliferation assay the three values per group are wells on a single plate, so they measure the variation between wells within one experiment, not the variation between experiments; the p values from Dunnett’s test should be read with that in mind. The A172 and A549 panels of Fig. 6 come from experiments performed on different days, and the dYB-1 experiment (Fig. 7B) is a third, separate experiment in which asRNA2 and dRNA2 did not differ from the scrambled-RNA control at day 3. The Western blot, RT-PCR and immunocytochemistry panels are single runs; the EGFR/ACTB ratios in Fig. 4 are single measurements rather than estimates of effect size. The ACTB signal in Fig. 5 fell alongside the EGFR signal, so that panel does not support a specific effect on EGFR mRNA, and repeating it as qPCR normalised to two or more reference genes is required before any claim about transcript abundance can be made.

Direct binding of YB-1 to the targeted *EGFR* sequence has not been tested; the assignment rests on SpliceAid prediction together with the behaviour of the dYB-1 decoy. RNA pulldown or RNA immunoprecipitation would be needed to establish it. Off-target effects of the oligonucleotides were not assessed, transfection efficiency was not measured, and all work is confined to a single glioblastoma line *in vitro*. U87MG and the temozolomide-resistant T98G line would extend the finding, and in vivo evaluation remains entirely open.

We post this preprint because the design strategy — using exon-definition logic and an RNA-binding protein prediction to choose an oligonucleotide target — may be useful to others working on *EGFR* in glioblastoma, and because the initial observations are reproducible enough in our hands to be worth recording. They are not sufficient to establish the mechanism we propose.

## Declarations

### Funding

This work was supported by the practical training budget of the Faculty of Pharmaceutical Sciences, Fukuoka University.

### Competing interests

The authors declare no competing interests.

### Author contributions

Conceptualization: K.O. Methodology: K.O. Investigation: Y.N. and K.O. Formal analysis: K.O. Visualization: K.O. Writing — original draft: K.O. Funding acquisition, supervision and project administration: K.O. Y.H., H.H., Y.M. and M.M. advised on study design and interpretation. Writing — review and editing: all authors. All authors read and approved the final manuscript.

### Data availability

All data generated in this study are presented in the figures.

## Notes

### Competing Interest Statement

The authors have declared no competing interest.

## References

1. National Cancer Center Japan, Center for Rare Cancers. Brain tumours. 2025.

2. Japan Society of Clinical Oncology. Clinical practice guidelines for brain tumours, 2nd ed. 2019.

3. Stewart LA. Chemotherapy in adult high-grade glioma: a systematic review and meta-analysis of individual patient data from 12 randomised trials. Lancet. 2002;359:1011–18.

4. Stupp R, Mason WP, van den Bent MJ, et al. Radiotherapy plus concomitant and adjuvant temozolomide for glioblastoma. N Engl J Med. 2005;352:987–96.

5. National Cancer Center Japan, Cancer Information Service. Treatment of glioma. 2023.

6. Japan Society for Neuro-Oncology. Glioblastoma guideline.

7. Louis DN, Perry A, Wesseling P, et al. The 2021 WHO Classification of Tumors of the Central Nervous System: a summary. Neuro Oncol. 2021;23:1231–51.

8. [Reference to be replaced with a primary source for the 35% EGFR amplification figure.]

9. Fenstermaker RA, Ciesielski MJ, Castiglia GJ. Tandem duplication of the epidermal growth factor receptor tyrosine kinase and calcium internalization domains in A-172 glioma cells. Oncogene. 1998;16:3435–43.

10. Ohe K, Maeda A. RNA processing. In: Jikken Igaku series, RNA. 2003:39–48.

11. Wang JZ, Zhu H, You P, et al. Upregulated YB-1 protein promotes glioblastoma growth through a YB1/CCT4/mLST8/mTOR pathway. J Clin Invest. 2022;132(8).

12. Zhou L, Lim QE, Wan G, Too HP. Normalization with genes encoding ribosomal proteins but not GAPDH provides an accurate quantification of gene expressions in neuronal differentiation of PC12 cells. BMC Genomics. 2010;11:75.

13. Dinh NTM, Nguyen TM, Park MK, Lee CH. Y-box binding protein 1: unraveling the multifaceted role in cancer development and therapeutic potential. Int J Mol Sci. 2024;25(2).

14. Evdokimova V, Tognon C, Ng T, et al. Translational activation of snail1 and other developmentally regulated transcription factors by YB-1 promotes an epithelial-mesenchymal transition. Cancer Cell. 2009;15:402–15.

15. Gong H, Gao S, Yu C, et al. Effect and mechanism of YB-1 knockdown on glioma cell growth, migration, and apoptosis. Acta Biochim Biophys Sin. 2020;52:168–79.

16. Zheng J, Dong W, Zhang J, Li G, Gong H. YB-1, a new biomarker of glioma progression, is associated with the prognosis of glioma patients. Acta Biochim Biophys Sin. 2016;48:318–25.

17. Bjorland LS, Fluge O, Gilje B, Mahesparan R, Farbu E. Treatment approach and survival from glioblastoma: results from a population-based retrospective cohort study from Western Norway. BMJ Open. 2021;11(3):e043208.

18. Yin Q, Zheng M, Luo Q, et al. YB-1 as an oncoprotein: functions, regulation, post-translational modifications, and targeted therapy. Cells. 2022;11:1217.

19. Bargou RC, Jürchott K, Wagener C, et al. Nuclear localization and increased levels of transcription factor YB-1 in primary human breast cancers are associated with intrinsic MDR1 gene expression. Nat Med. 1997;3:447–50.

20. Alkrekshi A, Wang W, Rana PS, Markovic V, Sossey-Alaoui K. A comprehensive review of the functions of YB-1 in cancer stemness, metastasis and drug resistance. Cell Signal. 2021;85:110073.

